# Human brain cortical oxygen consumption is positively related to soma surface area density

**DOI:** 10.64898/2026.09.29.755273

**Authors:** Manuela Carriero, Alessandra Caporale, Davide Di Censo, Ian Driver, Elizabeth J. Fear, Hannah Chandler, Fabrizio Fasano, Domenico Zacà, Emma Biondetti, Carolyn McNabb, Michael Germuska, Derek Jones, Marco Palombo, Richard G. Wise, Antonio M. Chiarelli

## Abstract

Evidence from in-silico, ex vivo and invasive animal studies suggested the function of neural tissue to be inextricably linked to its structural and microstructural organization. However, this link has never been assessed non-invasively in humans. Here, we established a simple quantitative framework linking cerebral oxygen metabolism to gray matter microstructural features in healthy humans using advanced non-invasive MRI methods. The cerebral metabolic rate of oxygen consumption (CMRO₂) was quantified with calibrated fMRI, whereas microstructural properties were estimated from diffusion MRI data fitted to the Soma and Neurite Density Imaging (SANDI) model, yielding soma radius (*R_soma_*) and soma volume fraction (*f_soma_*) metrics, from which we derived maps of soma surface area density (SAD) and soma numerical density (ND) under the assumption of a spherical soma. CMRO₂ was positively associated with SAD (r = 0.29, p < 0.01), *f_soma_* (r = 0.28, p < 0.05) and ND (r = 0.25, p < 0.05), and negatively associated with *R_soma_* (r = −0.23, p < 0.05) across cortical brain regions. These relationships are consistent with predictions from experimental and theoretical studies, providing the first quantitative evidence that cerebral metabolic demand is coupled to microstructure in humans. The positive association of energy consumption with SAD is consistent with the notion that both the number of synapses (energy-demanding structures), and the energy required to maintain a membrane potential, scale with soma surface area. Our findings establish a non-invasive MRI framework for investigating the microstructural determinants of brain energy metabolism in health and potentially in disease.

## Introduction

The microstructural features of brain tissue are central to its computational function^1–4^. Neurons composed of the soma, or cell body, axon and dendrites form the key computational unit at the cellular level. The electrochemical activity of these structures is one of the most energy-demanding processes in the human body, primarily relying on oxidative metabolism, with 90–95% of the brain’s ATP produced through oxidative phosphorylation. The brain requires a constant stream of glucose and oxygen, delivered through arterial blood, and accounts for approximately 20% of cardiac output despite constituting only 2% of adult body mass^5^. Furthermore, the majority of the brain’s energy expenditure occurs within the grey matter (GM). Although grey matter constitutes only about 40% of the total brain volume, it accounts for approximately 75–80% of the brain’s overall energy consumption. In contrast, white matter (WM), which comprises roughly 60% of the brain volume, consumes only about 20–25% of the total energy budget^6,7^.

The energy-consuming processes of neurons in GM can be divided into two main categories: i) signalling-related electrochemical processes, which account for roughly 75–80% of the cell’s energy budget — these include neurotransmitter cycling at synapses (∼43%), action potential propagation and the restoration of membrane potential after action potentials (∼17%), maintaining the resting potential of the cell membrane (∼15%); and ii) general housekeeping processes inside the cellular volume, including organelle transport, DNA repair, and protein synthesis, which account for the remaining 20–25%^8^. Conversely, in white matter (WM), glial cells and astrocytes are predominantly engaged in non-signalling maintenance functions, such as structural support and myelination. These housekeeping processes account for approximately 80% of the total WM energy expenditure, whereas signalling-related activities consume a substantially smaller proportion, estimated at around 20%^9^.

Relationships between energy consumption and microstructural characteristics in GM have been explored through *in silico* modelling^2^ and invasive experiments in animals^10^.

Principles of neural information theory establish that brain energy consumption and microstructure are not separate issues. Indeed, as formalized by Barlow^11^ and developed further by Stone^2^, Shannon’s information theory reveals that the brain is not merely a biological system but an engineered one, optimized to process information with maximal metabolic efficiency. By applying these mathematical principles in the visual system, it is possible to predict microstructural features such as the axon diameter^2^.

On the other hand, brain metabolic demands have been compared with brain microstructural characteristics across different mammalian species (mouse, rat, squirrel, macaque, baboon and human)^10^, where glucose utilization was measured using the [^14^C]2DG technique^12^ and the similar [^18^F]fluoro-2-deoxy-D-glucose^13^ method, and neuronal density was instead estimated using the isotropic fractionator technique^14^, which requires ex-vivo samples. When comparing across species, these studies found that total glucose use increased with the number of neurons. An other research study^15^ found the rate of glucose consumption to be correlated with synaptic density during mammalian brain development, with metabolic demand scaling proportionally to synapse number across species (rat, cat, monkey and human). They obtained these results by combining radioactive imaging data with data available on literature, mainly based on stereology or immunohistochemistry. Furthermore, bottom-up modelling based on anatomical and physiological data from rodents partitioned grey matter energy consumption among specific cellular compartments, including synapses, dendrites, axons, and glia^8^ and identified synaptic transmission as one of the dominant contributors to the brain’s energy expenditure^7^.

A recent study^4^ in humans quantified the association between MRI-derived Cerebral Blood Flow (CBF) and cellular density, i.e. cell-body staining intensity (CSI), using a new metric called CBF-CSI similarity index (CCSI). They found a region-specific alignment of CCSI, which is consistent with the varying metabolic demands and functional specialization of different cortical areas. Notably, this new metric was found to be related with mitochondrial respiratory capacity obtained through MitoBrainMap^16^. In addition, in this study, they also found CBF to be associated with the expression of genes related to the ion transmembrane transport across cell membrane. This suggests a link between brain energy demands and action potentials and synaptic transmission processes.

Finally, there is experimental evidence that a larger soma surface area can support a larger number of dendrites and synapses. By combining anatomical tracing and 3D digital reconstruction, comparative studies have reported that human pyramidal neurons from the temporal cortex are larger and more structurally complex than those in the primary sensory cortex, bearing a higher density of spines and synapses^17^.

Moreover, integrating spatial transcriptomics and in vivo (PET-FDG) neuroimaging, it was shown that primary motor cortex made of large pyramidal neurons (Betz cells) which generate efferent signals, demands substantial energy to support long-range axonal projections^18^. In addition to this, the neuronal cellular membrane may be modelled as a capacitor, owing to the ionic charge distribution and consequent potential difference between the intra- and extracellular compartments. The membrane capacitance, and thus the number of ions requiring energy to cross the membrane and maintain or restore a membrane potential, is proportional to the surface area of the capacitor^19,20^.

Considering the scientific evidence reported so far, we hypothesize that signalling-related electrochemical processes and therefore also energy consumption should primarily scale with neuronal soma surface area.

Larger neurons are likely to consume more energy, since a larger surface area implies more synapses and a larger membrane capacitance^20^. However, energy consumption is normally quantified per unit volume of tissue, or per unit mass, the two being equivalent under the assumption of uniform tissue density. Therefore, we expect the cell surface area per unit volume, or soma surface area density (SAD), to determine the energy consumption for populations of neurons. This leads us to predict a greater energy consumption, per unit volume of tissue, for smaller neurons, compared to larger neurons, as they deliver a larger surface area per unit volume of tissue for a given volume fraction occupancy.

Testing these hypotheses in humans requires simultaneous in vivo mapping of brain tissue energetics and microstructure. Recent developments in MRI permit to non-invasively investigate human tissue energetics in connection with microstructural features in vivo. Under the assumption of primarily oxidative metabolism, calibrated fMRI can map energy consumption in GM by quantifying the cerebral metabolic rate of oxygen (CMRO2). We recently demonstrated such an approach that can map CMRO₂ through BOLD and arterial spin labelling (ASL) recordings during a hypercapnic breath-hold task, by employing biophysical models of blood-oxygen-level-dependent (BOLD) signal and oxygen diffusion from capillaries to mitochondria^21–24^. Moreover, human brain tissue microstructure can now be studied using diffusion-weighted imaging (DWI) MRI. High b-value DWI, exploiting strong field gradients, permits quantitative assessment of microstructural tissue features; for example, the Soma And Neurite Density Imaging (SANDI)^25^ model estimates some cellular geometrical characteristics in GM.

The aim of the present study was to test a simple model linking brain energy consumption, characterised by CMRO_2_ assessed with breath-hold calibrated fMRI, to microstructural features revealed by DWI in cerebral GM. We primarily test the hypothesis, in healthy volunteers, that CMRO_2_ was positively associated with the soma surface area density (SAD), a structural metric evaluated for the first time in this study and derived from SANDI. Furthermore, we examine the dependence of CMRO_2_ on other microstructural features yielded by SANDI, the soma radius and the tissue volume fraction occupied by somas, together with an additional derived metric, namely the number of somas per unit volume, which we term the soma numerical density (ND). The associations between tissue energetics and structural features were assessed both across cortical regions and across subjects. These analyses may elucidate the importance of SAD in explaining brain energy consumption and identify which other cellular characteristics derived from DWI may drive variation in brain oxygen metabolism.

## Results

### Quantifying the Soma Surface Area Density (SAD) and Numerical Density (ND) *ND is consistent with literature*

We analysed data from 29 healthy subjects from the Welsh Advanced Neuroimaging Database (WAND)^26^, which includes maps of *CMRO*_2_ from the calibrated-fMRI method^24^ and microstructural parametric maps of the SANDI model from Diffusion Weighted Imaging (DWI)^25^.

SANDI models the diffusion weighted MRI signal in a voxel as a sum of three non-exchanging compartments (soma, neurite and extracellular space). Unlike earlier GM models^27,28^, it explicitly represents cell bodies (somas) as spheres thereby enabling estimation of the relaxation-weighted signal fraction from the soma compartment, *f_soma_*. Under the assumption of negligible relaxation-weighting, *f_soma_* represents the tissue volume fraction occupied by soma, termed the soma volume fraction. Moreover, SANDI estimates the “apparent” soma radius, *R_soma_*, which is the expected value of the distribution of soma sizes within the voxel. Please refer to the <u>Materials & Methods</u> section for details of the flow-diffusion model of oxygen transport used for *CMRO*_2_ estimation and of the SANDI model used for *R_soma_* and *f_soma_* estimation. Fig. 1 shows medians across subjects of *CMRO*_2_, *R_soma_* and *f_soma_* (respectively Fig. 1a, Fig. 1b and 1c) maps in MNI space.

**Fig. 1.**
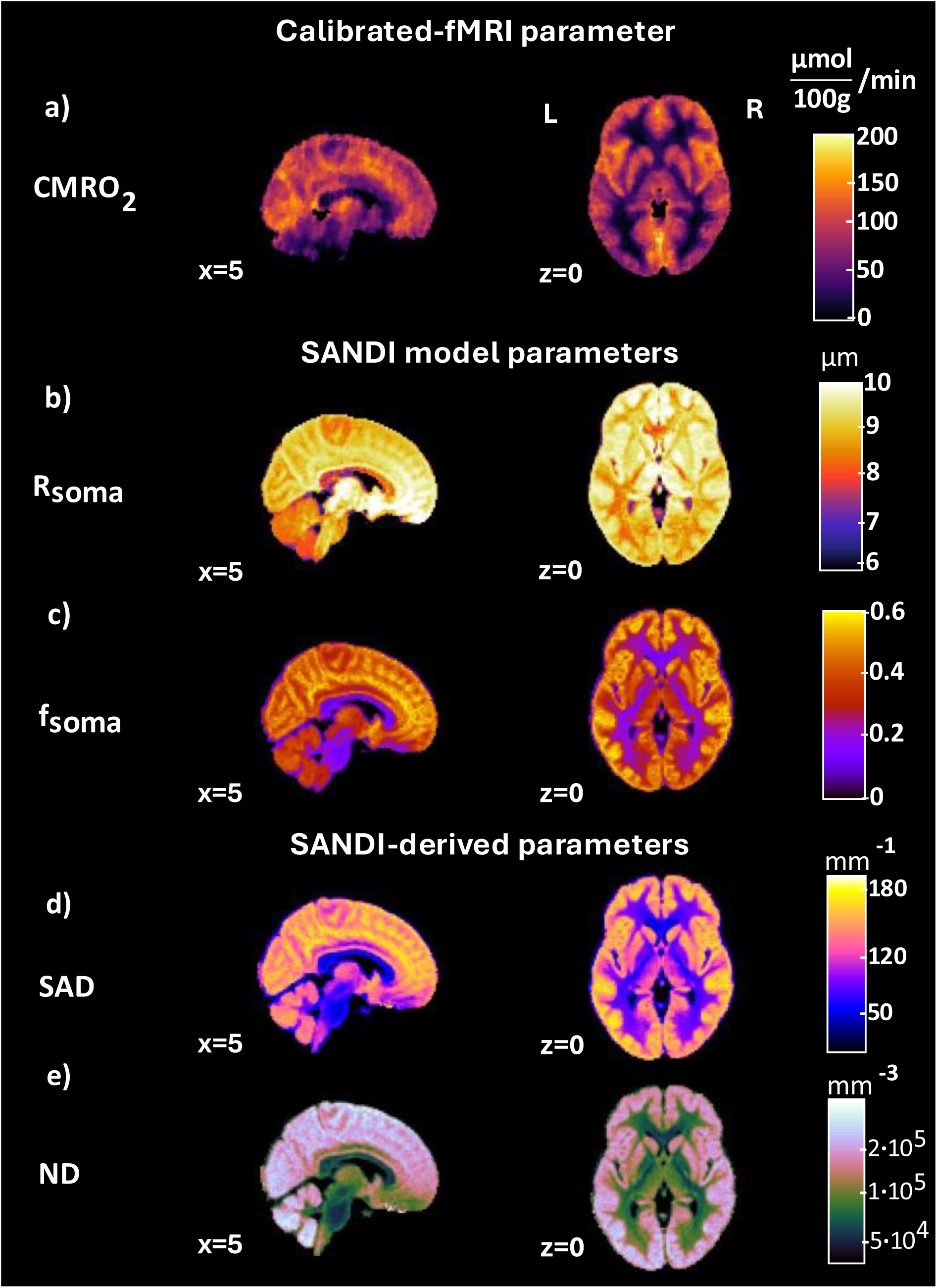
Parametric maps. Average maps of Cerebral Metabolic Rate of Oxygen Consumption, *CMRO*2 (a), soma radius, *Rsoma* (b), soma volume density, *fsoma* (c), soma Surface Area Density, SAD (d) and soma Numerical Density, ND (e), in MNI space, resolution 2×2×2 *mm*3, obtained as medians across 29 subjects. The coordinate origin (x=*z*=0) represents the midline of the brain in the left-right direction and in the superior-inferior direction.

Based on *R_soma_* and *f_soma_* we derived the following parameters: the soma surface area density (SAD), which is the surface area of cells per unit volume of tissue; and the soma numerical density (ND), which is the number of cells per unit volume of tissue. Assuming somas to be spheres and that relaxation-weighting effects on *f_soma_* are negligible, SAD and ND are equal to 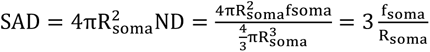, where the surface area and the volume of a single soma are S=4πR_soma_^2^ and 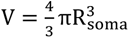 respectively (for fuller details of the model please refer to the <u>Materials & Methods</u> section). Fig. 1d and 1e show the medians across subjects in MNI space of the newly derived parametric maps, SAD and ND, respectively.

The order of magnitude of the ND map ∼ 10^4^ ÷ 10^5^ per cubic millimeter is consistent with histological studies results: Shapson-Coe and colleagues^29^ show that a cubic millimeter of human temporal cortex contains about 57,000 cells through electron microscopy reconstruction; a recent study derived 24,000 neurons/mm^3^ through a computational model of human cortical microcircuit^30^.

Furthermore, both the sagittal and axial view of the ND map in Figure 1e show a decrease of numerical cellular density from posterior to anterior regions, which is also observed in Ding et al.^31^ where digital Nissl-stained images of coronal human brain sections of the left hemisphere depict a higher cellular density in the posterior sections than in the anterior sections. Furthermore, a study from seven primate species have explicitly detected an anterior-to-posterior gradient in neuron density and number across the isocortex^32^, through the isotropic fractionator method^14^ and light microscopy. In such study, the authors present their findings as a fundamental feature of primate brain organization and they argue that the gradient should be present in the human brain, that is the largest primate brain. The sagittal view of Fig.1e shows also a greater intensity in the cerebellum with respect to the cortex, which is a well-established relationship in literature^14,33^.

No representative histological maps were found in the literature for the SAD parameter.

### Correlating CMRO_2_ with microstructure: between-regions analysis

#### Considering inter-regional variation, CMRO_2_ scales linearly with soma surface area density, soma volume fraction and soma numerical density (SAD, f_soma_ and ND)

We extracted the medians of the parameters in each ROI of the Automated Anatomical Labelling (AAL) atlas^34^ cortical regions and studied the spatial association of CMRO_2_ with microstructural parameters (i) for the ROI-wise median values computed across subjects and (ii) for each subject.

For both analyses we fitted a general linear model (GLM) using Equations (10)-(13) where CMRO_2_ and microstructural parameters were used as independent and dependent variables, respectively, as reported in <u>Materials & Methods</u>. The partial volume estimates of white matter (WM) and cerebrospinal fluid (CSF) were added as covariates to account for confounding effects on parameter estimation of mixed tissue types in each region.

Because individual imaging voxels typically contain contributions from multiple tissue types, GM, WM, and CSF, parameter estimates derived from such voxels are susceptible to non-GM signal contamination. Specifically, CSF and WM partial volume effects have been shown to attenuate functional signal estimates^35–37^ while artificially inflating diffusivity-based microstructural metrics^38^. This bias may therefore propagate to our derived biophysical parameters, namely CMRO₂ and SANDI metrics^39–41^. By modelling their linear effects as covariates, we can reduce the estimation error introduced by this partial volume confounds.

Fig. 2 shows the scatterplots depicting the spatial associations of regional median values of CMRO_2_ with regional median values of microstructural parameters where WM and CSF partial volume effects have been regressed out (median ROI-wise values across subjects were considered). All variables were standardized (z-scored).

**Fig. 2.**
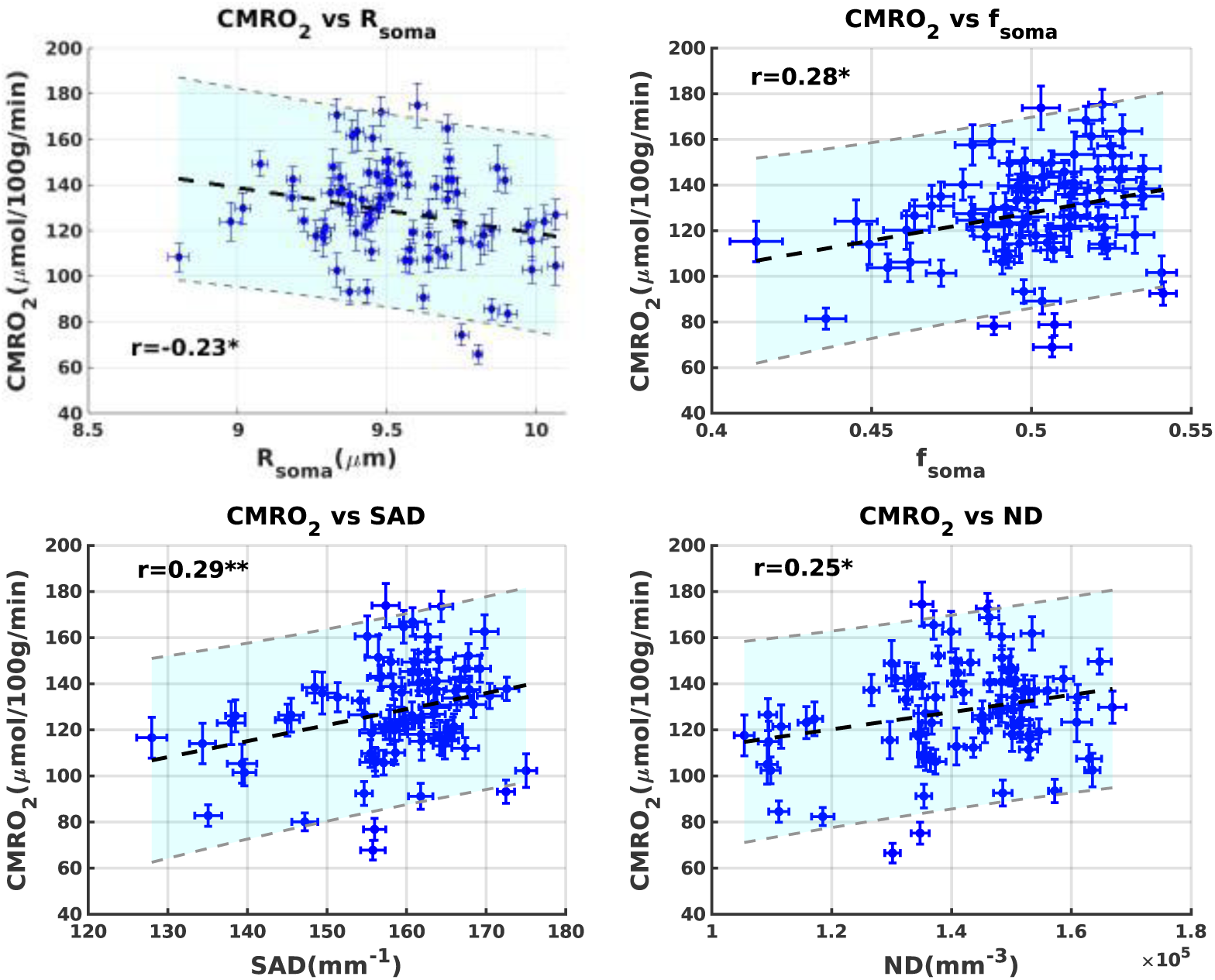
**Global association between *CMRO*_2_ and microstructural parameters across brain regions.** Scatterplots of spatial associations of *CMRO*_2_ with *R_soma_*, *f_soma_*, SAD and ND where WM and CSF partial volume effects have been regressed out and median values across subjects for each ROI were considered.

Associations between the CMRO_2_ and all the types of soma densities considered *f_soma_* , SAD and ND were positive and significant (Pearson correlation coefficients were respectively equal to *r* = 0.28, *p* < 0.05, *r* = 0.29, *p* < 0.01, *r* = 0.25, *p* < 0.01 ); a significant association, but with a negative sign, was found between *CMRO*_2_ and the soma radius *R_soma_* (*r* = −0.23, *p* < 0.05).

Fig. 3 shows the distribution (across subjects) of the regression coefficients, where regression coefficients evaluated, for each subject, the spatial association of *CMRO*_2_ with *R_soma_* (*α*_1_), *f_soma_* (*β*_1_), SAD (*γ*_1_) and ND (*δ*_1_). To facilitate interpretability and comparability of regression coefficients, all variables were z-scored prior to GLM analyses.

**Fig. 3.**
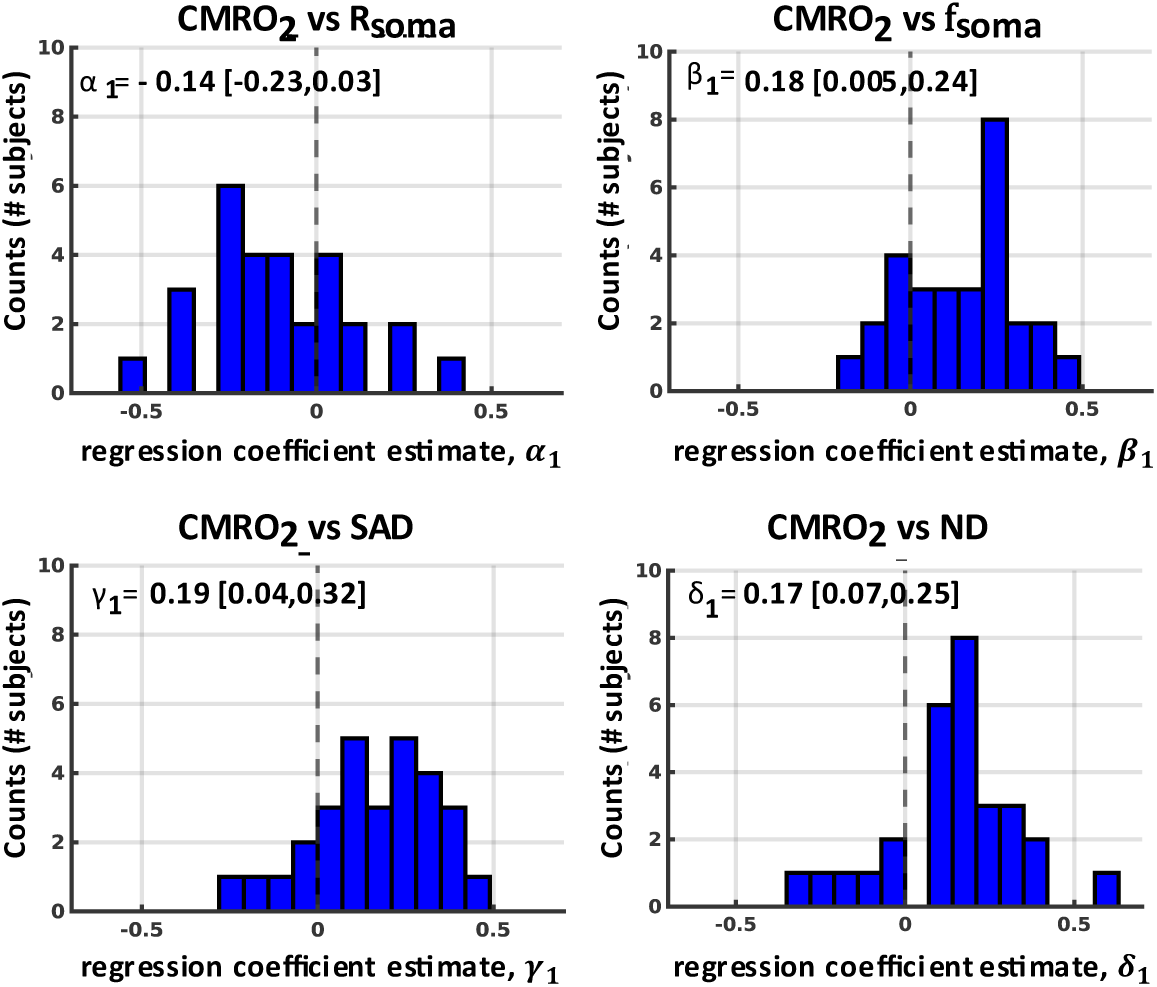
**Subject-wise association between *CMRO*_2_ and microstructural parameters across brain regions.** Distributions of regression coefficients between subjects, evaluating the across-ROIs spatial association of *CMRO*_2_ with *R_soma_*, *α*_1_, as in Eq. 9, *wit*ℎ *f_soma_* , *β*_1_, as in Eq. 10, with SAD, *γ*_1_, as in Eq. 11, and with ND, *δ*_1_, as in Eq. 12, where WM and CSF partial volume effects have been regressed out. Median and IQR (Q1–Q3) are shown in the upper-left corner.

For all regression coefficients, the mean of the distribution was significantly different from zero (p<0.05 for *α*_1_ distribution and p<10^−3^ for *β*_1_, *γ*_1_, *δ*_1_ distributions) with median and interquartile range (IQR[Q_1_,Q_3_]) respectively equal to *α*_1_ = −0.14 [0.23,0.03], *β*_1_ = 0.18 ± [0.005,0.24] , *γ*_1_ = 0.19 ± [0.04,0.32] and *δ*_1_ = 0.17 [0.07,0.25].

Each point represents a brain region. The bold black dashed line represents the linear regression line. The cyan shaded area, delimited by the grey dash lines, represents the 95% confidence intervals. * *p* < 0.05, ** *p* < 0.01.

### Correlating CMRO_2_ with microstructure: between-subjects analysis

#### Considering variation between subjects, CMRO_2_ is exclusively associated with soma surface area density (SAD) and soma volume fraction (f_soma_)

We considered two types of analyses which exploit between-subjects variability of the metrics evaluated, a global analysis focused on median values across the grey matter, and a regional analysis considering median values within multiple regions of interest (ROIs) derived from the AAL atlas^34^.

As in the previous section, we exploited GLM models with Equations (10)-(13) in the Materials & Methods section, adding WM and CSF partial volumes as covariates.

Fig. 4 shows the median grey matter *CMRO*_2_, having regressed out the effects of the covariates WM and CSF partial volumes, as a function of the four microstructural parameters considered: *R_soma_*, *f_soma_*, SAD and ND. The corresponding Pearson correlation coefficients are also reported in the plots.

**Fig. 4.**
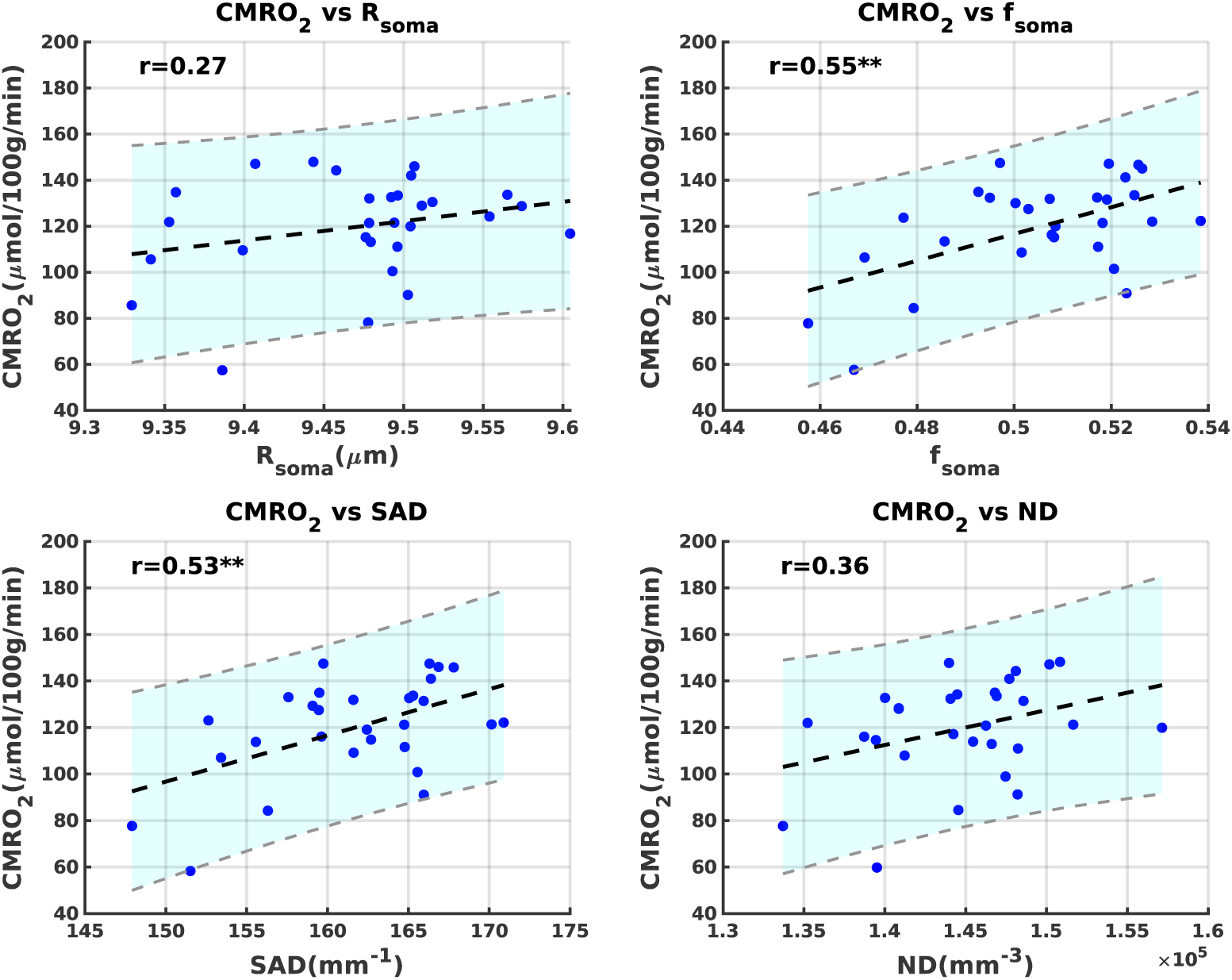
**Global association between *CMRO*_2_ and microstructural parameters across subjects.** Scatterplots of grey matter median values of *CMRO*_2_ vs *R_soma_*, *f_soma_*, SAD and ND where WM and CSF partial volume effects have been regressed out. Each point represents a subject. The black bold, dashed line represents the linear regression line. The cyan shaded area, delimited by the grey dash lines, represents the 95% confidence intervals. ** *p* < 0.01.

The correlation coefficient was significant between *CMRO*_2_ and *SAD* with *r* = 0.53, *p* < 0.01 and between *CMRO*_2_ and *f_soma_* with *r* = 0.55, *p* < 0.01; the correlation was not significant between *CMRO*_2_ and *R_soma_*, and *CMRO*_2_ and *ND*.

Fig. 5 shows maps of the correlation coefficient obtained between CMRO_2_ and the microstructural parameters when considering GM segmented into ROIs, with only significant regions being displayed (p<0.05, FDR-corrected). CMRO_2_ was positively associated across subjects with all four microstructural parameters considered. However, compared with *R_soma_* and ND, SAD and *f_soma_* display a higher number of regions with a significant positive association with CMRO_2_.

**Fig. 5.**
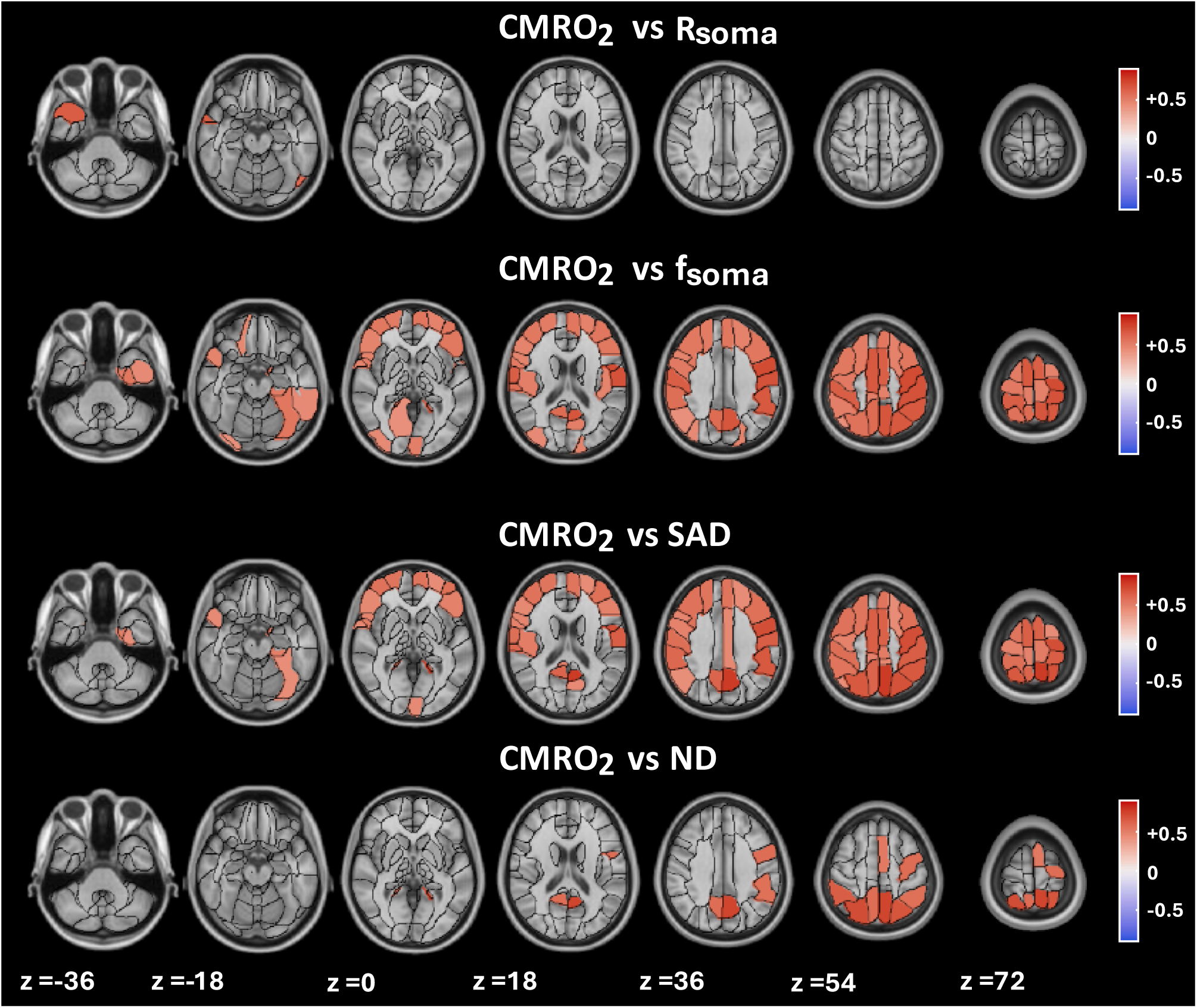
**For each ROI, association between *CMRO*_2_ and microstructural parameters across subjects.** Axial slices showing ROI-based Pearson correlation coefficient for *CMRO*_2_ vs *R_soma_*, *f_soma_* , SAD and ND, where WM and CSF partial volume effects have been regressed out. Benjamini–Hochberg (BH) FDR-corrected, *p* < 0.05.

## Discussion

We performed a study that combined breath-hold calibrated fMRI with high-gradient diffusion-weighted imaging in healthy volunteers to map, in grey matter, the cerebral metabolic rate of oxygen consumption (CMRO₂) and tissue microstructural properties, respectively. We derived specific microstructural properties related to neuronal somas, including the soma surface area density (SAD). Consistent with our hypothesis, when accounting statistically for partial-volume contamination of CSF and WM, we observed that CMRO₂ was positively associated with SAD, considering both between-region and between-subject variability. As with SAD, the association between CMRO₂ and the soma volume fraction, *f_soma_*, was positive both between regions and between subjects.

The results are consistent with the notion that the principal energy-demanding processes scale with the sum of the soma surface areas present in an image voxel or SAD as we have defined it. To date, several *in silico* and *in vivo* studies suggest that the cell surface hosts the most energy demanding structures and processes. Inter-neuronal signal transmission relies on neurotransmitters whose cycling happens at the presynaptic terminals^42^ and accounts for a significant portion of energy consumption^43^. Hence, the number of synapses strongly influences the amount of energy required by a cell. Studies^17^ suggest that the surface area of the soma determines the number of synapses a cell can harbour^44^. Moreover, we expect the surface area of somas present to determine the energy required to maintain and restore the resting membrane potential. Owing to its distribution of charges, the neuronal cell acts as a capacitor^45^, storing charges to keep the resting membrane potential at around −70mV. Because the capacitance is proportional to the capacitor surface area, a greater surface areas should consume more energy, since a higher quantity of charge is needed to maintain a given potential difference^20^. These considerations apply to the single cell but when evaluating energy consumption in a given volume of tissue the sum of cellular surface area within that volume (surface area density, SAD) is relevant. For a given tissue volume fraction occupied by soma (f_soma_), SAD increases as R_soma_ decreases.

Consistently with this reasoning and the observed positive association between CMRO_2_ and SAD, we identified a negative association between CMRO_2_ and the *R_soma_* across ROIs, both when considering across-subjects ROIs medians and when considering the relationships for each subject.

However, when considering the global GM value, the between-subject association of CMRO_2_ with soma radius was non-significant (Fig. 4), likely due to the smaller variability of soma radius values, which ranges only from 9.3 to 9.6 μm in the case of subjects’ global GM median values but from 8.8 to 10.1 μm in the case of ROIs’ medians across subjects. Furthermore, across subjects, only two regions had a significant correlation coefficient surviving multiple comparison correction (Fig. 5). However, the number of regions with a significant association was much lower than the number of regions with a significant association between CMRO_2_ and all other types of density parameters (SAD, *f_soma_*, and ND).

As with SAD, *CMRO*_2_ was found to be positively associated with soma volume fraction, *f_soma_*, both considering variability across subjects and across space. This result was expected both from a biological point of view, as the cell volume supports several energy-consuming processes (for instance, protein-synthesis processes that occur inside a cell^46^), and from a physical point of view, as the volume is positively related to the surface area.

Another expected significant association was that between CMRO_2_ and the soma numerical density ND, which was positive and significant across ROIs, both considering the global analysis and the analysis for each subject. This is also confirmed in a study across species through ex-vivo samples, in which the total glucose use was found to increase with the number of neurons^10^. However, the trend was still positive but not significant across subjects when considering GM medians samples (Fig. 4). The number of ROIs with significant positive association across subjects was smaller than the ones found between CMRO_2_ and *f_soma_* and SAD, but still higher than the ones between CMRO_2_ and *R_soma_* (Fig. 5).

Although the results were consistent across ROIs when considering the global associations and the associations for each subject, only the positive associations between CMRO_2_ and surface area density (SAD) and volume density (*f_soma_*) held across subjects. Overall, microstructural parameters exhibited less variability across subjects than across regions, which may explain why the associations between CMRO_2_ and the soma radius *R_soma_* and between CMRO_2_ and the soma numerical density (ND), were non-significant. On the contrary, as the relationships between CMRO_2_ and SAD and *f_soma_* were still observed when considering variability between subjects, this suggests that SAD and *f_soma_* are the quantities that best correlate with CMRO_2_.

The microstructural parameters derived from the SANDI model, SAD and ND, represent a non-invasive alternative to histology. To date, diffusion MRI has largely been limited to providing information about the proportion of the total MRI signal originating from specific compartments which is usually taken to represent the fraction of the volume occupied by those compartments. For instance, the SANDI model parameter *f_soma_* is the relaxation-weighted volume fraction occupied by somas. In white matter, too, we can estimate the axonal volume fraction, from models such as NODDI^27^, which estimates the proportion of the MRI signal originating from water molecules within axons. The present study, by contrast, exploiting the ability of the SANDI model to provide estimates of brain-cell geometry (i.e. the soma radius), moves beyond these “volumetric parameters” by calculating two specific densities, the SAD and ND parameters, using Eqs. (6) and (8), respectively.

Notably, the order of magnitude (∼10^4^ ÷ 10^5^ number of cells per cubic millimetre) of the ND map agrees with literature values using histological^29^ and a computational model^30^. Additionally, the ND map shows a decrease of its intensity from posterior to anterior regions. This is the same trend as is observed in the Nissl-stained images of the cellular-resolution atlas^31^ and in the brain of seven primate species^32^ though the isotropic fractionator method^14^ and light microscopy performed on ex-vivo samples of brain tissues. This finding provides qualitative validation of the derived ND parametric map and represents a step forward in the evolution of methods for counting brain cells. In contrast, SAD lacks histological validation, reflecting the novelty of this parameter across fields.

In evaluating these relationships, we considered the cerebrospinal fluid and white matter partial volume as potential confounding factors and we, therefore, included them in the regression model. SANDI microstructural parameters in GM are affected by partial volumes errors from other tissues present in the voxel ^39–41^. We expect estimates of soma radius and volume density parameter to increase with increasing CSF contamination and decrease with WM contamination. We expect contamination effects for the *CMRO*_2_ parameter as well, that is derived from BOLD and ASL measurements on which the partial volume effect has been observed in several studies^35–37^, in this case leading to a decreased functional signal where CSF and WM contamination is present.

To minimize partial volume contamination, our analyses were restricted to the cortical regions of the AAL atlas. Subcortical structures, such as the thalamus, hippocampus, and amygdala, are located in close proximity to WM and are therefore inherently more susceptible to partial volume effects arising from neighbouring tissue compartments^47^. Additionally, recent studies have reported lower inter-subject reproducibility of SANDI-derived parameters in subcortical regions, likely reflecting the reduced signal-to-noise ratio (SNR) in these structures^48^.

The role of the 3T Connectom MRI system is important in obtaining interpretable microstructural data. *R_soma_* and *f_soma_* parametric maps from the SANDI model were derived from diffusion weighted imaging (DWI) data acquired using the Connectom MRI scanner that, thanks to the use of the stronger gradients, decreasing the diffusion times and the degree of *T*_2_-weighting as well as increasing the signal-to-noise ratio, allows for reliable estimation of SANDI model parameters and minimizes the influence of relaxation-weightings on volume fraction estimates^25^.

### Limitations

The soma surface area density (SAD) and the soma numerical density (ND) were obtained by combining two SANDI model parameters: the soma volume fraction, *f_soma_* , and soma radius, *R_soma_*. This approach assumes a spherical somas whereas, in reality, they have different and irregular shapes^49^. Moreover, the SANDI model soma compartment represents the cell body of any brain cell type, which inherently includes contributions from both neurons and non-neuronal cells such as astrocytes, microglia, and oligodendrocytes. Therefore, the microstructural metrics derived in this study do not purely reflect neuronal properties, despite our hypothesis being based on neuronal properties. However, given that our analysis was focused on grey matter, where neuronal cell bodies are abundant, we interpret these parameters as predominantly reflecting neuronal microstructure, with the understanding that they may also be influenced by the non-neuronal population. Future studies are needed to further disentangle the relative contributions of these different cell types.

In addition to the geometrical and cell-type assumptions, interpretation of SANDI parameters assumes negligible relaxation weighting effects on the different compartment fractions. However, different tissue compartments (soma, neurite and extracellular space) exhibit different T_2_ and T_1_ relaxation times^25,50,51^.

Consequently, the estimated compartmental signal fractions are likely to be relaxation-weighted, potentially reflecting a combination of volume fractions and compartment-specific relaxation properties rather than pure volume fractions alone. These effects are minimized using the Connectome scanner which, thanks to the strong gradients, can decrease the T_2_ weighting. The present work may be developed further by the use of advanced multi-TE acquisitions to quantify separate intra-neurite and intra-soma T₂ relaxation times^51^ and accounting for T_1_ properties of the tissue.

Moreover, all analyses were performed assuming linearity, both for assessing associations and for accounting for, by linear regression, the CSF and WM partial volume contaminations.

Regarding the biological interpretation of our findings, we acknowledge that multiple factors may underlie the observed relationships. Among these are electrophysiological effects that could not be quantified in the present study. For instance, *CMRO*_2_, as a measure of metabolic consumption rate, may be influenced by the temporal dynamics of electrochemical activity, such as neuronal firing rate. Firing rate, in turn, may either depend on microstructural features, thereby mediating the association between *CMRO*_2_ and microstructural parameters, or be independent of microstructure. Additionally, our observed energy–microstructure relationships could be driven by molecular and genetic factors, as suggested by a recent study demonstrating that both the coupling strength between CBF and cytoarchitecture, as well as CBF itself, are associated with specific molecular, cellular, and transcriptomic profiles^4^. These considerations highlight the need for future multimodal investigations to further elucidate the mechanisms underlying the associations identified in this work.

MRI technical limits have also led us to focus on cortical GM. Functional MRI studies for CMRO_2_ quantification in WM are nowadays still limited by the low SNR^52^ and the SANDI model is less suited to WM due to low *f_soma_*^53^. Overcoming these technical limits, would lead us to have a complete picture of the identified relationships in all brain tissues.

Finally, we hypothesized that the positive association of CMRO₂ with the soma surface area density (SAD) is primarily driven by the association of SAD with the number of synapses, as shown by histological studies focused on specific brain regions^17,44^. However, this inference could be further explored with MRI, using DWI protocols and models able to probe even smaller spatial scales and thereby estimate the number of synapses. DWI models currently under development aim to quantify the number of dendritic spines on a cell^54^, but their translation to in vivo experiments is still under investigation. Future advances in DWI models may enable us to further develop and refine our brain energy–microstructure model.

## Conclusion

We present a novel model that explains the dependence of energy consumption on specific microstructural metrics in healthy human grey matter, using non-invasive, in-vivo MRI methods. According to our findings, CMRO_2_ is associated with the total surface area of cells per unit volume of tissue (the soma surface area density, SAD) and with the soma volume fraction (*f_soma_*). Between brain regions, CMRO_2_ is found to be positively associated with the soma numerical density (ND) and negatively associated with the soma radius (*R_soma_*), supporting the notion that CMRO_2_ scales with the surface area present in the tissue.

Finally, it is worth noting that the newly derived density maps extracted expanding the DWI SANDI model, namely SAD and ND, may serve as new quantitative parameters to be used for the non-invasive characterization of brain microstructure via DWI.

## Materials & Methods

### Data collection

Data from twenty-nine healthy volunteers (age: 24.4±5.9 years; F/M: 12/17) of the Welsh Advanced Neuroimaging Database (WAND)^26^ were considered for the study. We included subjects who underwent both breath-hold calibrated fMRI and diffusion weighted image acquisition on the 3T Connectom MRI system, to have *CMRO*_2_ and SANDI microstructural parameter maps for each subject. The models and the parameter estimation techniques are summarized in the following sections. The study was approved by the Cardiff University School of Psychology Research Ethics Committee (EC.18.08.14.5332RA3).

### MRI data

Breath-hold calibrated fMRI and DWI data were acquired using two different 3T scanners, MAGNETOM Prisma and Connectom (Siemens Healthineers, Forchheim, Germany) respectively at Cardiff University Brain Research Imaging Centre (CUBRIC), Cardiff, UK. In both cases, a 32-channel receive-only head coil from the same vendor was used.

Calibrated fMRI was performed using an in-house dual-excitation^55^ (DEXI) pCASL sequence with two inversion pulses for background suppression, which allows simultaneous acquisition of BOLD and ASL cerebral blood flow images. The labelling duration and the Post Label Delay (PLD) were both set equal to 1.5 s, 3-fold in-plane GRAPPA undersampling was used with *TE*_1_ = 10 *ms* and *TE*_2_ = 30 *ms*. A TR of 4.4 s was used to acquire 15 slices of 2D EPI, with in-plane resolution 3.4 × 3.4 mm and slice thickness 6 mm (33% slice gap).

The 10 minute DEXI protocol was run with a breath-hold task consisted of 10 repeats of a 20 s end-expiration breath-hold, after every one of which was a recovery period. In between breath-holds, participants were visually cued to pace their breathing at 3 s inhale, 3 s exhale. Visual prompts were presented on a video screen at the centre of the participant’s field of view.

To enable CBF quantification and susceptibility distortion corrections, a pair of proton density images (M_0_s) were acquired with opposite phase encoding directions (A≫P and P≫A) with TR 6000 ms, TE=10 ms and no tagging. EPI parameters and resolution were chosen to match those of the DEXI acquisition.

The Diffusion Weighted MRI dataset was acquired through a single-shot spin-echo, echo-planar imaging sequence. Data were acquired in both anterior to posterior (A≫P) and posterior to anterior (P≫A) phase-encoding directions. A≫P data comprised two shells (b = 200 *s*/*mm*^2^ and b = 500 *s*/*mm*^2^) with 20 diffusion encoding directions uniformly distributed (for full details of the directions, see McNabb et al.^26^), one shell (b = 1200 *s*/*mm*^2^) with 30 directions and three shells (b = 2400 *s*/*mm*^2^, 4000 *s*/*mm*^2^and 6000 *s*/*mm*^2^) with 61 directions, in addition to two initial non-diffusion-weighted (b = 0 *s*/*mm*^2^) images and 11 non-diffusion-weighted images dispersed throughout the acquisition (33rd volume and every 20th volume thereafter). P≫A data comprised two initial non-diffusion-weighted images, one shell of 30 directions at b = 1200 *s*/*mm*^2^ and a final non-diffusion-weighted image. Data acquisition details for all b-values were as follows: TR 3000 ms, TE 59 ms, FOV 220 × 220 × 132 *mm*^3^, voxel size 2 × 2 × 2 mm, with 2-fold in-plane undersampling. Diffusion gradient duration (δ) and separation (Δ) were 7 ms and 24 ms, respectively.

T1-weighted anatomical data were acquired using the 3 T Prisma MRI scanner and exploited for segmentation and registration purposes. The MRI sequence was MPRAGE with the following acquisition parameters: TR 2250 ms, TE 3.06 ms, flip angle 9°, FOV 256 × 288 × 176 *mm*^3^, voxel size 1 × 1 × 1 mm, TI 850 ms, 2-fold in-plane undersampling and phase-encoding direction A≫P.

### MRI Analysis

This section describes the preprocessing pipelines for the fMRI and DWI datasets and the methods for model parameter estimation.

DEXI-pCASL data consisted of two time series, one for *TE*_1_ (ASL images) and one for *TE*_2_ (BOLD images) which were rigid-body motion corrected (6 degrees of freedom) using MCFLIRT^56^. ASL and BOLD images were corrected for susceptibility distortions using the pair of M0 images acquired with reversed phase encoding directions, through FSL ApplyTopup ^57^ function. Perfusion-weighted and BOLD-weighted time series were derived via surround tag-control subtraction and surround averaging of the *TE*_1_ and *TE*_2_ time series, respectively.

ASL control-tag difference perfusion data (Δ*S*) was normalized with respect to the M_0_ and converted to Cerebral Blood Flow (CBF) quantitative units of ml/100g/min through the pCASL single compartment kinetic model of labelled spins and voxel-wise signal normalization^58^:

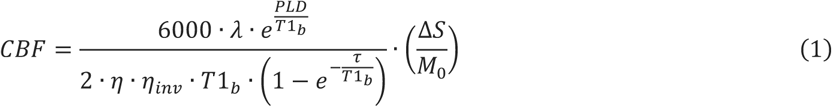

Where k is the water partition coefficient (k=0.9mL/g), *T*1*_b_* is the T1 relaxation constant of blood, *η* is the tagging inversion efficiency (*η*=0.85) and *η_inv_* is a scaling factor to account for the reduction in tagging efficiency due to background suppression (*η_inv_*=0.88)^59^. The *T*1*_b_* was calculated from the arterial oxygen saturation of hemoglobin, SaO2 and partial pressure of oxygen in arteries,PaO_2_, using the experimental relation presented in David T. Pilkinton et al.,2012 ^60^. PaO_2_ was assumed to be equal to P_ET_O_2_ extracted from O_2_ recordings, whereas SaO_2_ was calculated from PaO_2_ using the Hill equation^22^.

The mean timeseries from ASL and BOLD voxels within the GM was used to create a physiological regressor representing the global breath-holding stimulus, with the GM selected considering a GM mask created through the mean perfusion signal thresholded between the 85^th^ and 99^th^ percentiles^23^. Regression against ASL and BOLD timeseries was then used to determine the resting Cerebral Blood Flow, CBF_0_, the fractional change in CBF, and the BOLD signal change in response to hypercapnia induced by breath-holding^61,62^.

CMRO_2_ was evaluated via the Fick Principle:

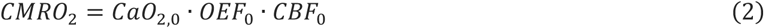

where OEF_0_ is the oxygen extraction fraction and CaO_2,0_ is the arterial concentration of oxygen at rest.

CaO_2,0_ was evaluated from end-tidal O_20_ traces. OEF_0_ was instead estimated using the approach described in detail in Chiarelli et al., 2022^22^. The method infers OEF from the maximum BOLD modulation (M) derived from comparing BOLD and CBF fractional changes following hypercapnia via the equation^22,23^:

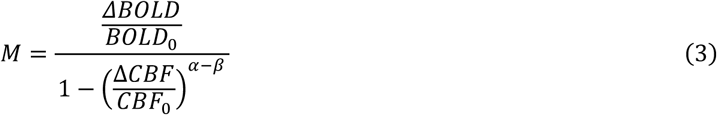

Using the central volume theorem^21^ and the flow-diffusion model of oxygen transport^22^, the equation relating M to OEF_0_ becomes:

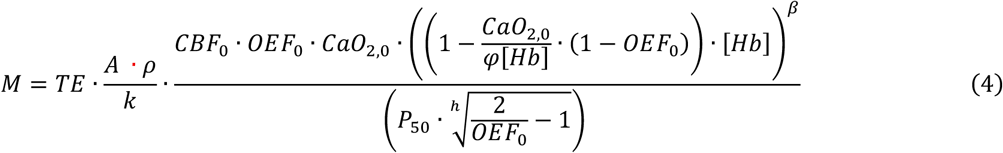

where k is the effective oxygen permeability of brain tissue, *β* and A are field strength and vessel geometry dependent constants, *φ* is the oxygen binding capacity of hemoglobin; [Hb] is the concentration of hemoglobin in blood. P_50_ is the oxygen partial pressure when half of Hb is saturated and h is the Hill constant (h=2.8). *TE* is the sequence echo time. *ρ* is a proportionality constant, used to relate the capillary volume *CBV_cap_* to the broader venous compartment *CBV_v_* , that is *CBV_v_* = *ρCBV_cap_*.

The inversion recovery sequence was used to quantify the T1 of venous blood^63^ for estimation of systemic [Hb]. We assigned a value of 8.8 s^-1^g^-β^dl^β^/(μmol/mmHg/ml/min) to the term (A_GE_·ρ)/k, matching our previously established *in-vivo* estimates^22^. *β* = 1.3 at 3T, *φ* = 1.34mL/g. Generally P_50_ ≈ 26mmHg; P_50_ can be inferred from a measure of end-tidal partial pressure of carbon dioxide, P_ET_CO_2_.

By comparing M from the flow–diffusion model (Equation 4) with the calibrated fMRI expression for M under hypercapnia^22^, OEF₀ can be determined by minimizing the difference between the two estimates.

DWI data were pre-processed via signal drift correction with SOLID^64^, denoising through MPPCA and Gibbs ringing correction (MRTRIX3^65^), susceptibility and eddy-current-induced distortions corrections (FSL^57,66,67^) and gradient non-uniformity correction using DISCO^68^.

We then applied the novel Soma and Neurite Density Imaging (SANDI)^25^ model to estimate microstructural parameters. Previous biophysical diffusion models assumed water molecules to diffuse in two non-exchanging compartments: the intracellular and the extracellular spaces. SANDI models the intracellular compartment as comprising two sub-compartments: the neurites and the cell body (i.e. the soma). This microstructural model assumes non-exchanging compartments for short diffusion times *t_d_* (*t_d_* ≤ 20*ms*).

The model estimates orientation-independent microstructural features by considering the average of the signals acquired along uniformly distributed directions. The resulting signal takes the form:

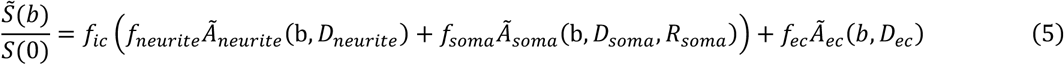

Where *S̃*(***b***) is the average of the directions *S*(0) and b is the diffusion weighting factor; *f_ic_* and *f_ec_* are the intra-cellular and extra-cellular relative signal fractions satisfying the condition *f_ic_* + *f_ec_* = 1; *f_neurite_* and *f_soma_* are the intracellular compartments signal fractions, namely the neurite and soma relative signal fractions satisfying the condition *f_neurite_* + *f_soma_* = 1; *Ã _neurite_*, *Ã _soma_* and *Ã _ec_* are the direction-averaged normalised signals associated respectively with the neurite, soma and the extracellular compartment; *D_neurite_*, *D_soma_*, *D_ec_* are the diffusion constants of the intracellular compartments, neurite and soma, and extracellular compartment, respectively; *R_soma_* is the soma radius. The total free parameters to be determined from the direction-averaged data are thus six: *f_neurite_* , *f_ec_* , *D_neurite_*, *D_soma_*, *D_ec_* and *R_soma_*. Given the difficulty in disentangling *D_soma_* from *R_soma_*, *D_soma_* value is fixed equal to 3*μm*^2^/*ms*.^25^

SANDI model parameter maps were retrieved through model inversion based on a machine-learning framework applied to synthetic data. The approach involved generation of synthetic signals with Rician and Gaussian noise as input for a random forest which was trained to invert the SANDI model. The software used is the Matlab SANDI Toolbox available in GitHub. The procedure is explained and validated elsewhere ^25^. The Rician noise was added considering the Marchenko-Pastur Principal Component Analysis (MPPCA)^69,70^ noise map calculated on images acquired with b-values = 0,200, 500,1200,2400 *s*/*mm*^2^^71^; the Gaussian noise was instead extracted from the residual of the spherical harmonics decomposition of the signal acquired in-vivo^72^.

### Definition of new microstructural parameters

The SANDI model assumes somas to be spheres and it yields an estimation of the average soma radius within a voxel, *R_soma_*, and the soma signal fraction, *f_soma_*, representing the relaxation-weighted volume fraction occupied by soma, Equation 5. Assuming *f_soma_* equal to the soma volume fraction (i.e., negligible effects of relaxation-weighting), we can define the soma surface area density, *SAD*, as the ratio of the estimated surface area of a single soma (S=4*πR*^2^ ) to its volume (V=^4^ *πR*^3^ ) weighted by the soma volume fraction:

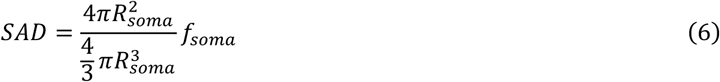

Simplifying Equation (6), *SAD* can be calculated simply as:

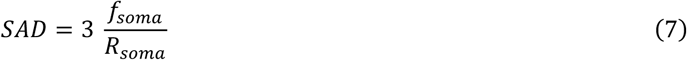

Equation (6) can be rewritten as:

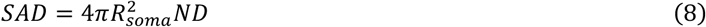

where ND is the ratio of soma volume fraction to the volume of a single soma, i.e.

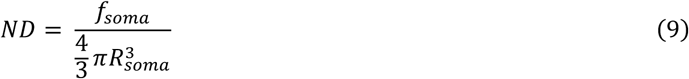

representing the number of cells per unit volume.

### General Linear Models

Relationships between metabolic and microstructural parameters were evaluated using General Linear Models^73^ via two distinct approaches: one focusing on inter-regional variability and the other on inter-subject variability. In the former, regional medians were treated as independent samples, both within each subject and taking the medians across subjects, to explore spatial variability across brain areas. Furthermore, for the inter-subject variability analysis, we considered the global medians of parameters across the whole of grey matter; we also performed a parcellation-based analysis, analysing the median values of each region across the subjects. We opted for a ROI-based median analysis rather than a voxel-wise approach to enhance the signal-to-noise ratio (SNR) and minimize statistical dependence between samples.

The Automated Anatomical Labelling (AAL) Atlas^34^, MNI space, voxel size 2 × 2 × 2 *mm*^3^, containing 170 ROIs, was used to parcellate the brain into regions.

Data analysis was performed in subject space. To map the AAL atlas regions onto the functional and diffusion spaces, we used *AntsApplyTransforms*^74^ with *MultiLabel* interpolation. For each subject, the warping was achieved by concatenating the inverse of two sequential transformations: (i) the alignment of the reference M_0_ (or b_0_ for diffusion) to the T1-weighted MPRAGE and (ii) the registration of the MPRAGE to MNI152 template. Specifically, the former (i) was executed via a rigid-body transformation, while the latter (ii) employed a rigid-body followed by an affine transformation and, finally, a non-linear transformation. Both (i) and (ii) were implemented through *AntsRegistration*^74^.

The new derived microstructural maps, SAD and ND, were calculated through Equations (7) and (8).

GM, WM and CSF partial volume estimate (PVE) maps were calculated through FAST^75^ applied to MPRAGE images which were brain extracted through BET^76^. Each voxel in these maps contains a value between 0 and 1, representing the estimated proportion of the corresponding tissue class. These maps were registered to the functional/diffusion space through the inverse of the transformation (i) relative to M_0_/b_0_, using *AntsApplyTransforms* with *NearestNeighbor* interpolation.

A GM partial volume estimate threshold of 0.5 (i.e., 50%) was then applied to all parametric maps to select GM tissue.

As both CMRO_2_ and SANDI model parameters may be influenced by partial volume contamination from WM and CSF^35–41^, we included the regional average values of partial volume estimates (or whole GM averages, depending on the analysis) as covariates. We considered the WM and CSF maps in the in the diffusion space, as it was the space with higher resolution.

The GLM equations modelling the relationship between the CMRO_2_ and the microstructural parameters *r_s_* , *f_s_*,*SAD*, *ND*, for both subjects’ and regional variability analysis, were the following:

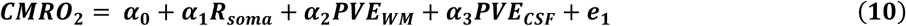

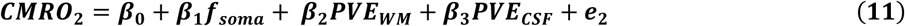

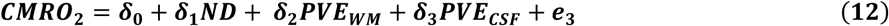

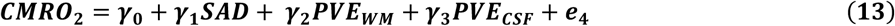

where *α*_0_, *β*_0_, *δ*_0_, *γ*_0_ are the intercept and *α_i_*, *β_i_*, *δ_i_*, *γ_i_* are the GLM parameter estimates, with *i* = 1, . . , *N*, with N equal to the number of covariates. *e*_1_, *e*_2_, *e*_3_, *e*_4_ are the random error terms.

Both for the inter-regional (medians across subjects) and the inter-subjects variability analysis, Pearson correlation coefficients^77^ were then computed after having regressed out the contribution of WM and CSF PVEs from the CMRO_2_ variable. The significance of the correlation was assessed using a t-test, as implemented in the Matlab’s *corrcoef* function. Only correlation coefficients with p < 0.05 were considered statistically significant.

For the inter-regional variability analysis, the regression coefficients *α*_1_, *β*_1_, *δ*_1_, *γ*_1_ were computed for each subject and the *α*_1_, *β*_1_, *δ*_1_, *γ*_1_ distributions were evaluated through one-sample t-test and by computing the median values and the interquartile (IQR) range. To facilitate interpretability and comparability of regression coefficients, all variables were z-scored prior to GLM analyses.

## Code availability

Python 3.11 was used to plot figure 1 and all plots and statistical analysis were performed using MATLAB version 9.13.0 (R2022b) (The MathWorks Inc., Natick, MA). All codes are available on GitHub (https://git.cardiff.ac.uk/cubric/wand, https://github.com/ManuelaCarriero/EMI_model).

## Acknowledgments

The WAND data were acquired at the UK National Facility for In Vivo MR Imaging of Human Tissue Microstructure funded by the EPSRC (grant EP/M029778/1), and The Wolfson Foundation, and supported by a Wellcome Trust Investigator Award (096646/Z/11/Z) and a Wellcome Trust Strategic Award (104943/Z/14/Z). The UK Medical Research Council (MR/M008932/1), the Welsh Government and Cardiff University provide funding support for the CUBRIC ultra-high field imaging facility. Moreover, AC was partially supported by the Fund for the Promotion and Development of Policies of the National Research Program, as per DM 737/2021 issued by the MUR. D.D.C. was supported by EU-NextGeneration EU-MUR, PNRR, Project Title: “VITALITY – Innovation, digitalization and sustainability for the diffused economy in Central Italy”. EJF was supported by EU-NextGenerationEU-MUR, National Plan for Recovery and Resilience (PNRR) and Projects of National Relevance (PRIN), project number: P2022ESHT4, project Title: “Advancing MRI biomarkers of brain tissue microstructure and energetics in Multiple Sclerosis.” Funding call No. 1409 of 14.09.2022, Concession decree No. 1367 of 01.09.2023 adopted by MUR, ERC Panel LS5 “Neuroscience and Disorders of the Nervous System”. CUP: D53D23019210001. R.G.W. was partially supported by EU-NextGeneration EU-MUR, Projects of National Relevance (PRIN), project Code 2022MHMSSJ, project Title: “Combining magnetic resonance imaging and optical spectroscopy to map microvascular function and oxygen metabolism in healthy and diseased brain”. CUP: D53C24004560006. A.M.C. was partially supported by EU-NextGeneration EU-MUR, Research National Program (PNR) and PRIN. Project Code: 2022BERM2F, Project Title: “Mapping Mitochondrial Function and Oxygen Metabolism in the Human Brain with Magnetic Resonance Imaging”. Funding call No.104 of 02.02.2022, Concession decree No. 1065 of 18.07.2023 adopted by MUR, ERC Panel LS7 “Prevention, Diagnosis and Treatment of Human Diseases”. CUP: D53D23013410001. In collaboration with Siemens Healthineers.

## Declaration of conpeting interests

Manuela Carriero received research support from Siemens Healthineers (Forchheim, Germany).

